# Revealing the relevant spatiotemporal scale underlying whole-brain dynamics

**DOI:** 10.1101/2020.09.12.277699

**Authors:** Xenia Kobeleva, Ane López-González, Morten L. Kringelbach, Gustavo Deco

## Abstract

The brain rapidly processes and adapts to new information by dynamically switching between activity in whole-brain functional networks. In this whole-brain modelling study we investigate the relevance of spatiotemporal scale in whole-brain functional networks. This is achieved through estimating brain parcellations at different spatial scales (100-900 regions) and time series at different temporal scales (from milliseconds to seconds) generated by a whole-brain model fitted to fMRI data. We quantify a fingerprint of healthy dynamics quantifying the richness of the dynamical repertoire at each spatiotemporal scale by computing the entropy of switching activity between whole-brain functional networks. The results show that the optimal relevant spatial scale is around 300 regions and a temporal scale of around 150 milliseconds. Overall, this study provides much needed evidence for the relevant spatiotemporal scales needed to make sense of neuroimaging data.

## Introduction

The brain can rapidly process and adapt to new information through the flexible transitioning between multiple states. Functional neuroimaging studies demonstrate how the macroscopic brain organization dynamically changes during these transitions of multiple functional states, even in the absence of an active task (Tang et al. 2012; Stitt et al. 2017; Liégeois et al. 2019). There has been convincing evidence that brain dynamics rest on the orchestrated activity of several networks of brain regions which transition in recurring patterns over time (Alexandrov 1999; Meer et al. 2020). These transitions between brain networks have been associated with to cognition and (ab)normal behaviour (Engel et al. 2001; Thompson et al. 2013; Vidaurre et al. 2017; Liégeois et al. 2019; Lurie et al. 2020; Yoo et al. 2020). However, a fundamental question remains, namly at which particular spatiotemporal scale the whole-brain functional networks are able to optimally transition.

The current body on research on spatiotemporal scales of the dynamical behaviour of whole-brain networks is limited, since empirical studies of different spatial and temporal scales are challenging. In human neuroimaging studies, spatiotemporal scales have a restricted range for each modality. The spatial resolution of fMRI is now down to less than a millimetre but it is not clear if this is the right scale for capturing the richness of information processing across the whole-brain. Similarly, the spatial resolution of MEG depends on the sensors and it has been shown that beamforming can only separate up to around 70 regions across the whole-brain with significant drop in signal in deeper regions. Even if the acquisition of whole-brain imaging is now around 0.7 seconds, the temporal resolution of fMRI is limited by the haemodynamics of the BOLD signal.

Dynamic whole-brain models offer an elegant opportunity to overcome the limitations of the restricted spatiotemporal scales in experimental research (Yuan et al. 2018; Deco et al. 2019; Cornblath et al. 2020). Using a whole-brain network model, in our previous research we were able to compare the complexity of dynamic switching behaviour of whole-brain networks across different time scales from milliseconds to seconds (Deco et al. 2019). In this study, we extend our previous work on different temporal scales (i.e. the temporal resolution) in a whole-brain network model (Deco et al. 2019) by adding a spatial dimension to the analysis (i.e. the number of regions) and explore the switching behaviour of networks across spatiotemporal scales (i.e. taking into account both spatial and temporal scales). By doing so, we attempt to answer the question at which spatiotemporal scale macroscopic whole-brain functional networks can provide optimal richness of repertoire. Thus, our study has implications for a better understanding of the dynamical reconfiguration of whole-brain functional networks over time. We aim at providing a quantification of how best to choose appropriate neuroimaging modalities and parcellation techniques when investigating the dynamics of whole-brain functional networks, keeping the balance between maximum information content and computational complexity of the analysis. In this study we focus on the dynamic behaviour of macroscopic, functional brain networks and use the simplest form to quantify the richness of the dynamical repertoire, using an entropy measure.

To achieve our goal, we explore the switching behaviour of whole-brain functional networks at spatial scales from 100 to 900 regions both in empirical time series extracted from resting-state fMRI with fixed temporal scales as well as in simulated time series with various temporal scales from milliseconds to seconds. We determine the relevant spatiotemporal scale by comparing the entropy of the switching activity. In information theory, entropy describes the level of variability of a given variable (Shannon 1948). By focusing on the behaviour of whole-brain networks, we focus on *relevant* information in brain dynamics and find the maximum of the entropy, which allows us to choose the most *optimal* spatiotemporal scale. In the discussion of our results, we derive recommendations for neuroimaging researchers, highlighting our finding that the relevant spatial scale for analyses of brain dynamics is around 300 regions and at an optimal temporal scale of around 150 milliseconds and thus contribute to an empirical basis of relevant parameters for studies of brain dynamics.

## Methods

We adapted the existing comparing different time scales (Deco et al. 2019) to incorporate different spatial scales. Images were created using Biorender, Inkscape, Connectome Workbench and the Matplotlib library within Python.

### Data acquisition and preprocessing

We used resting state functional MRI data from 100 unrelated subjects of the Human Connectome Project (HCP; Van Essen et al. 2013) with a mean age of 29.1 ± 3.7 years. The HCP study was approved by the local ethical committees and informed consent was obtained from all subjects. Six subjects were discarded as the resulting FC matrices consisted of at least one not available row at parcellations with more than 800 regions (due to the sparsity of the networks). We further chose one of the four available resting-state fMRI scans of about 15 minutes duration (TR of 0.72 sec). During fMRI acquisition, subjects were instructed to keep their eyes open while looking at a fixation cross. A full description of the imaging parameters and minimal preprocessing pipeline can be found in Glasser et al. (2013). In short, after correction for motion, gradient and susceptibility distortions the fMRI data was aligned to an anatomical image. The aligned functional image was then corrected for intensity bias, demeaned and projected to a common surface space, which resulted in a cifti-file. All fMRI data was filtered between 0.1 and 0.01 Hz to retain the relevant frequency range for further analyses of the BOLD signal. We obtain structural and functional matrices in different spatial scales using the Schaefer parcellation, which optimizes local gradient and global similarity measures of the fMRI signal in various spatial scales ranging from 100 to 900 regions (Schaefer et al. 2018). In both fMRI datasets time series were extracted with the help *Workbench Command* provided by the HCP.

To create a structural connectome as a basis for the whole-brain model, we generated a structural connectome depicting the number of fibers in the required spatial scales. We used the diffusion MRI dataset from the HCP database, that uses high-quality scanning protocols with an acquisition time of 89 minutes for each of the 32 participants, resulting in above-average normative diffusion MRI data. The data has already been preprocessed and made available to the public within the Lead-DBS software package (Setsompop et al. 2013; Horn et al. 2017). In brief, the data were processed using a generalized q-sampling imaging algorithm as implemented in *DSI studio* (http://dsi-studio.labsolver.org). The data were segmented and co-registered using *SPM 12*. Restricted by a coregistered white-matter mask, 200,000 fibers were sampled within each participant using a Gibbs’ tracking approach (Kreher et al. 2008) and normalized into MNI space via DARTEL transforms (Ashburner 2007; Horn and Blankenburg 2016). We used the standardized methods from *Lead-DBS toolbox* version 2.0 (Horn et al. 2018) to obtain structural connectomes for the same parcellation schemes as for the functional data, selecting tracts that both started and ended within the specified parcellation scheme.

### Whole-brain modeling using the DMF model

The use of fMRI signals would normally limit our study in the temporal dimension. To overcome this shortcoming, we use a whole-brain model which allows us simulate data in varying timescales from milliseconds to seconds, while a comparable structure of the signal. We create a dynamic mean field (DMF) model, which is conceptually based on interconnected regions containing excitatory and inhibitory neuronal pools (Deco et al. 2013).

A summary of the individual steps that were taken to create the model can be found in Figure 1. The model consists of a network of brain regions that emit spontaneous neuronal signals. The number of the brain regions is defined by the spatial scale. Each of these regions consists of excitatory (*E*) and inhibitory (*I*) neuronal pools that reciprocally influence each other locally within each region. We further assume that these regions interact via long-range connections, as given by the connection weights of the structural connectome (Deco et al. 2014).

**Figure 1.**
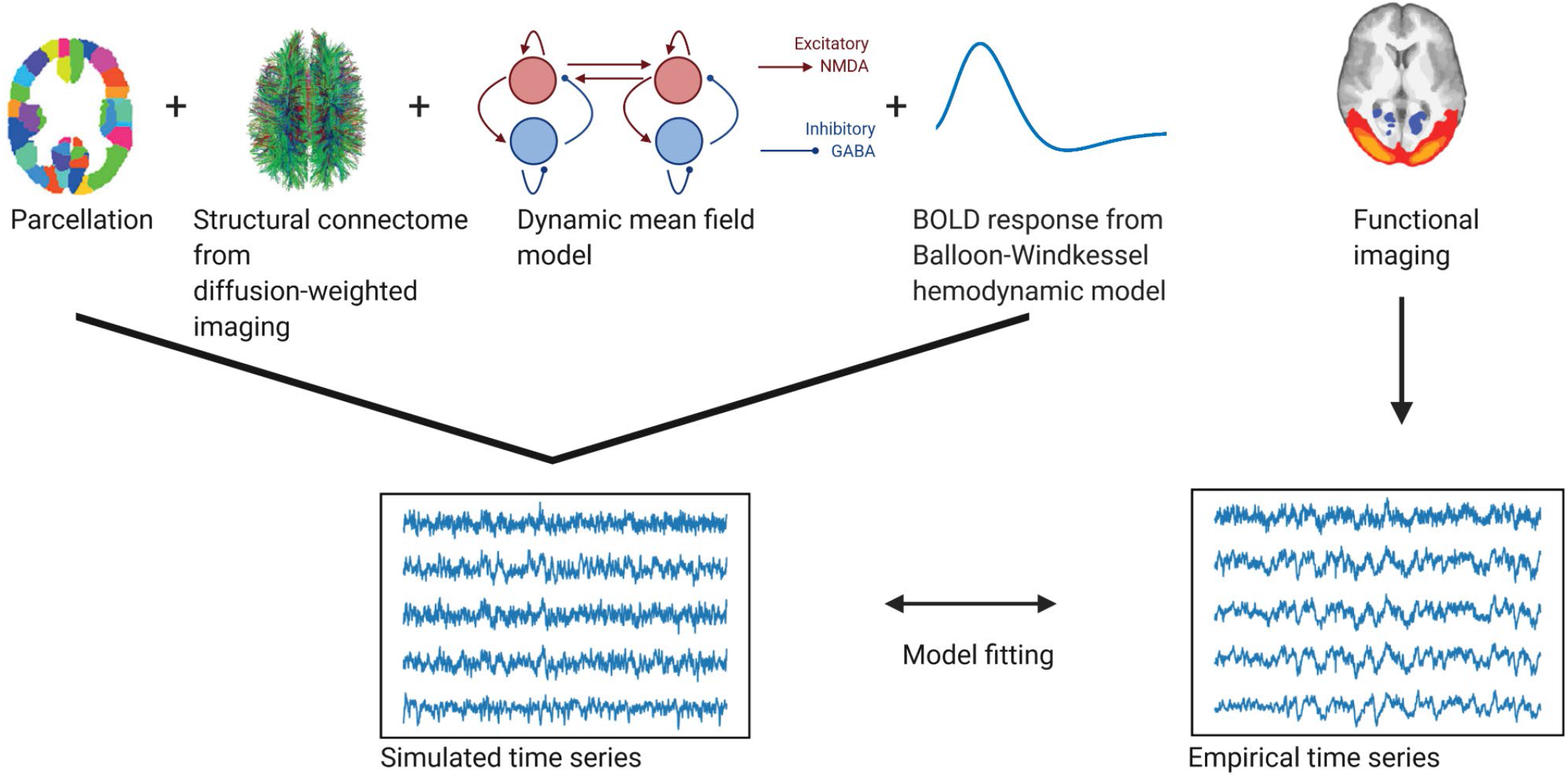
Whole-brain modeling steps to create simulated functional time series fitted to empirical BOLD data. Using a whole-brain network model such as the dynamic mean field model allows us to accurately create time series data at different temporal scales. Local dynamics of each region given by a *parcellation* are generated by a *dynamic mean field model* and coupled through the *structural connectome* (as provided by the numbers of fiber tracts estimated from diffusion-weighted imaging). To fit the resulting neuronal time series to the empirical BOLD time series, we employ a *Balloon-Windkessel hemodynamic model* to create simulated BOLD time series. The simulated time series are *fitted* to the empirical time series using metrics of metastability and phase similarity matrix distributions.

These assumptions are implemented through a modified DMF model based on the original reduction first proposed by Wong and Wang (2006). In the model used in this study, NMDA receptors mediate excitatory currents *I*^(*E*)^ and GABA-A receptors mediate inhibitory currents *I*^(*I*)^. Inhibitory sub-populations communicate reciprocally with excitatory sub-populations on a local level. Excitatory sub-populations are additionally linked to other excitatory sub-populations via long-range connections, representing the effect of NMDA receptors. These long-range connections are based on the number of fiber tracts given by the structural connectome (see description above). The connections are then tuned by a global scaling factor G that linearly scales all synaptic strengths.

The following set of coupled differential equations are used to create the DMF model:

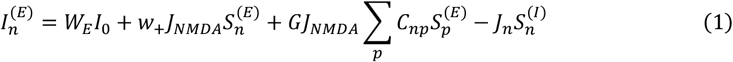

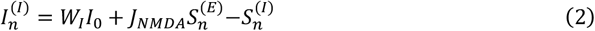

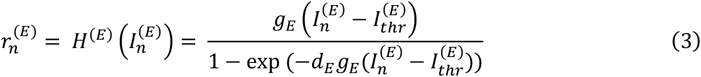

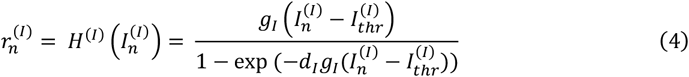

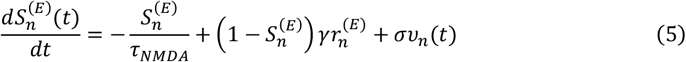

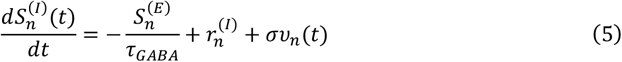

For each inhibitory (*I*) and excitatory (*E*) neuronal pool in every brain region *n*, the vector 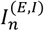 Represents the total input current (in nanoamperes), the vector stands for the firing rate (in 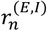 hertz) and the vector 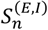 denotes the synaptic gating. The total input currents that are received by the neuronal pools are converted by the neuronal response functions *H*^(*E* ,*I*)^into firing rates 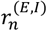. Here, the gain factors *g*_*E*_ = 310 nC^−1^ and *g*_*I*_ = 310 nC^−1^ are used to determine the slope of *H*. When the threshold currents of 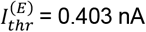 and 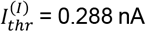 are reached,the firing rates increase linearly with the input currents. The shape of the curvature of H around *I*_*thr*_ is defined by the constants *d*_*E*_ = 0.16 and *d*_*i*_ = 0.087. The average synaptic gating of the excitatory pools 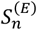 is controlled by the NMDA receptors with a decay time constant *τ*_*NMDA*_ =0.1 s and *γ*= 0.641(transformed into ms). The average synaptic gating of the inhibitory pools 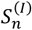 is controlled by the GABA receptors with a decay time constant *τ*_*GABA*_= 0.01 s (transformed into ms). All excitatory synaptic couplings are weighted by *J*_*NMDA*_ = 0.15 nA and the weight of the recurrent excitation *w*_+_= 1.4. The overall effective external input is *I*_0_ = 0.382 nA with *W*_*E*_ = 1 and *W*_*I*_ = 0.7. We add standard Gaussian noise *v*_*n*_ with an amplitude of *σ* = 0.01 nA. To mimic a resting state condition, the weight of feedback inhibition *J*_*n*_ is adjusted for each excitatory subpopulation to obtain a firing rate 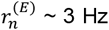. This was done using a regulatory mechanism called Feedback Inhibition Control, which was shown to mimic resting state activity better (Deco et al. 2014).

It is then possible to retrieve separate temporal scales from the simulated neuronal data by binning the time series. However, first the neuronal time series had to be fitted to the empirical BOLD time series (by adjusting *G*) to ensure a biologically plausible signal. Therefore, we transformed the neuronal signal from the model into a simulated BOLD signal and then compared the simulated and empirical signals (see below). We employed the Balloon-Windkessel hemodynamic model using all biophysical parameters as stated in (Stephan et al. 2007). The model is described by the following equations:

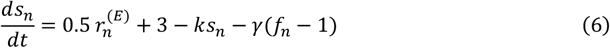

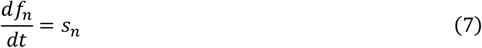

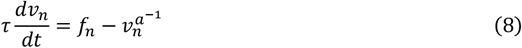

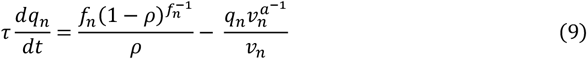

This model describes a vasodilatory signal *s*_*n*_ which is altered by autoregulatory feedback. Depending on *s*_*n*_, the blood flow *f*_*n*_ leads to changes of the deoxyhemoglobin content *q*_*n*_ and blood volume *v*_*n*_. *τ* is the time constant, *ρ* is the resting oxygen fraction and *a* represents the venous resistance. For each region *n* the BOLD signal *B*_*n*_ is a static nonlinear function of *q*_*n*_ and *v*_*n*_:

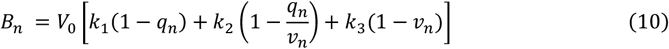

To focus on the functionally relevant frequency range, we band-pass filtered the simulated BOLD signals using the same filter as for the empirical data with a bandpass between 0.1 and 0.01 Hz (Achard et al. 2006; Glerean et al. 2012).

### Agreement between empirical and simulated data

To achieve biologically plausible signal statistics in the simulated time series at each scale, we performed the fitting to the empirical signals by adjusting G to have a maximal agreement in two different metrics: the metastability, and phase consistency matrices (see below). Each of these metrics represents different dynamical properties of the BOLD signal. Previous research has showed that these adding dynamical metrics such as metastability and phase consistency matrices are better at constraining dynamical working points of dynamical whole-brain models than using static metrics such as FC only (Deco et al. 2017, 2019; Saenger et al. 2017). These metrics were computed for each value of G (between 0 and 2.5 in steps of 0.025) in the simulated data and for the empirical data and compared as described below. Due to multiple spatial scales, the creation of the model was very compute-intensive, e.g. to replicate the time series of 10 subjects from the HCP dataset at a neuronal timescale using a parcellation of 400 regions with different G-values from 0 to 2.5 about 80-100 GB of RAM & 30 days of computation were required. Therefore, we restricted the simulations to 10 iterations, representing time series of a group of 10 subjects. To prove that our analyses were generalizable to a larger group of healthy subjects, we did 100 iterations of the model fitting to empirical time series of a group of 10 subjects from the HCP dataset, that were randomly selected at each iteration.

#### Dynamical measures used for the fitting

##### Metastability

The metastability represents the overall variability of oscillations (Wildie and Shanahan 2012; Deco et al. 2017). It is calculated as the standard deviation of the Kuramoto order parameter R(t) across time, which depicts the average phase φ*k*(*t*) in a given region *k* across *n* regions.

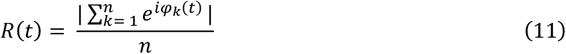

The phases were derived from the data by detrending the filtered fMRI time series and then applying the Hilbert transform. When *R* = 1 all phases are fully synchronized, while *R* = 0 indicates a complete desynchronization of all phases. We calculated the differences between the empirical and simulated metastability. This has been previously proven to be suitable to define the dynamical working point of dynamical whole-brain models (Deco et al. 2017; Saenger et al. 2017).

##### Phase consistency matrices

We calculated the phase coherence matrix by evaluating the instantaneous phase at each time point *t* of every region *j* and then computing the phase difference across all regions. We measured the similarity of these phase coherence matrices over *t* to create a *phase consistency matrix*. This resulted in a representation of spatiotemporal fluctuations of phases. To compare between empirical and simulated data, we calculated the Kolmogorov-Smirnov distance between the empirical and simulated distribution of the phase consistency matrices. The Kolmogorov-Smirnov distance quantifies the maximal difference between two distribution functions of two samples and is minimized by the optimal value of G (Saenger et al. 2017).

Furthermore, we checked whether we retrieved comparable numbers of functional networks in the empirical and simulated data (see Figure S1 in the supplementary data).

### Extraction of whole-brain functional networks using independent component analysis and calculation of entropy

The summary of the analytical steps can be seen in Figure 2. The simulated and empirical time series were available in different spatial scales. In the case of the simulated signal, we aimed to retrieve the simulated neuronal time series at separate temporal scales in the range of milliseconds to seconds (see Figure 2A). To do so, the simulated neuronal time series were binned by averaging the signals in windows of the width of the timescale, each time bin corresponding to a time point of the newly created time series. As this approach led to multiple fine-grained time series with a high computational cost of the analysis, we were only able to simulate the time series across all temporal scales up to a spatial scale of 400 regions. We created simulated time series at group level by performing 10 iterations (representing 10 subjects).

**Figure 2.**
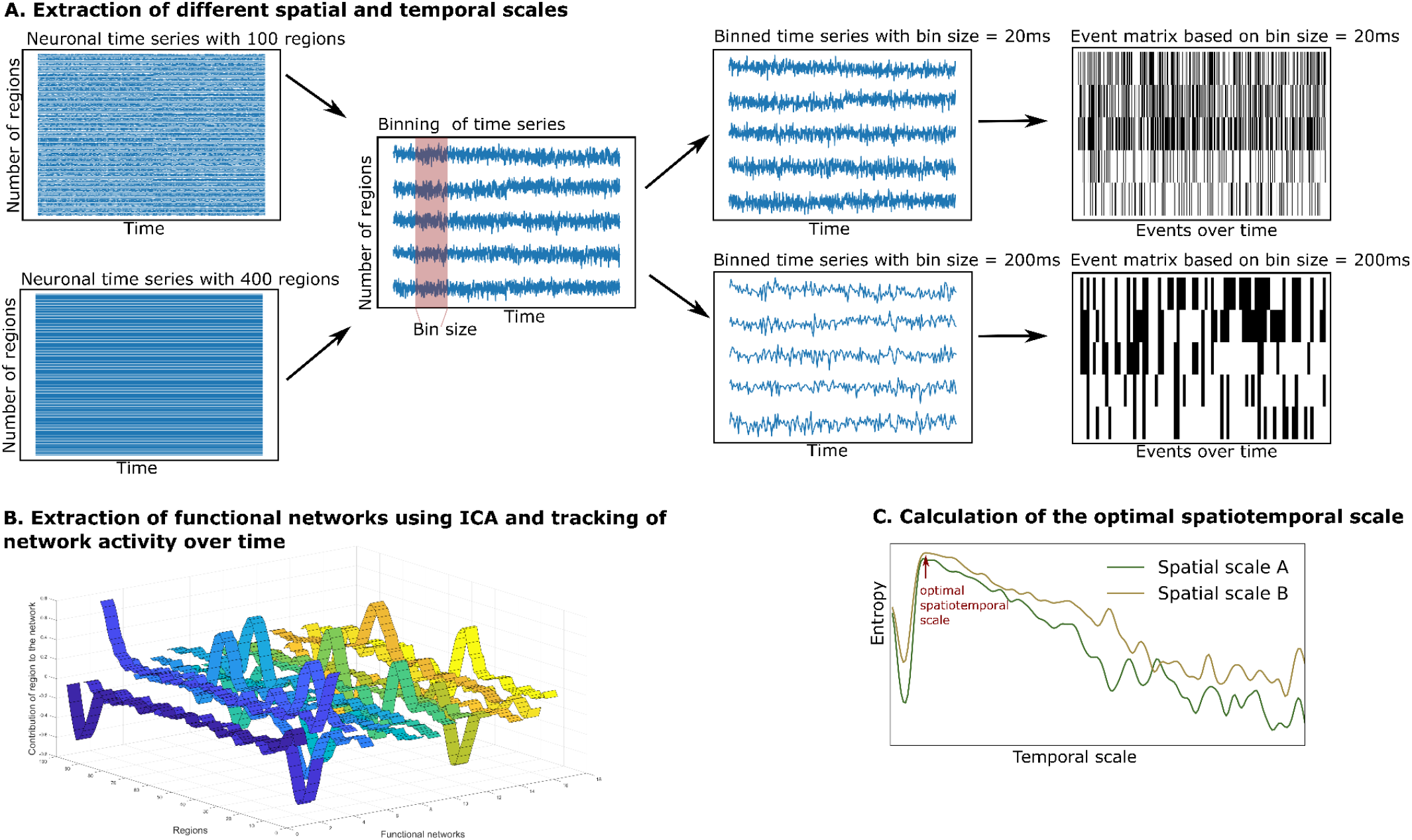
Extraction and tracking of whole-brain functional networks at different spatial and temporal scales using the whole-brain model. A. We simulate neuronal time series at different spatial scales (from 100 to 400 regions). We then create different bin sizes of the time series (using bins from 10ms to 3000ms), the bin size corresponds to the temporal scale. The binned time series are binarized using a point process paradigm, resulting in an event matrix. B. We extract whole-brain functional networks using independent component analysis, resulting in a network matrix (see ribbon plot) These networks are tracked over time by projecting the event matrix onto the networks, resulting in an activity matrix (not displayed). C. The richness of the switching between functional networks is estimated by calculating the entropy of their switching probability. The entropy is compared across spatial and temporal scales.

In the case of the empirical time series, we extracted a group of 10 subjects from the data by randomly selecting 10 subjects. We concatenated their time series to retrieve functional networks on a group level (using the same group size as in the simulation to ensure comparability). To make the analysis robust to interindividual variability, we repeated this process 100 times. The temporal scale of the empirical data was determined by the TR (HCP: 720 ms). Given only one temporal scale we were able to extract functional networks in a spatial scale from 100 to 900 regions.

In each temporal scale (given by the TR in the empirical data or the bin size in the simulated data), the time series were binarized using the point-process binarization algorithm for BOLD signals (Tagliazucchi et al. 2012). Here, the time series were normalized using a *z*-score transformation and depending on a threshold the time series were set to 0 or 1, resulting in an event matrix (see the right panel of Figure 2A). Next, the event matrix was normalized using z-score transformation, so that the event matrix in each brain region would have null mean and unitary variance. This procedure has been shown to be robust to threshold choices and is a classical method to reduce dimensionality of dynamical data (Tagliazucchi et al. 2012). We then continued the analysis with the normalized event matrix *e* (with the dimension: number of regions *i* x number of time points *b*).

To estimate the number of functional networks, we applied an adaptation of an eigenvalue analysis for assessing the statistical significance of resulting networks (Peyrache et al. 2010; Deco et al. 2019), as introduced by Lopes-dos-Santos, Ribeiro, and Tort (2013). This method finds the number of principal components within the event matrix that have significantly larger eigenvalues compared to a normal random matrix that follows a probability function, as specified in Marčenko and Pastur (1967). As can be seen in Figure 2B (left panel), after determining the number of functional networks, we extracted these functional networks by applying an independent component analysis to the event matrix *e*. This procedure resulted in a resulting in a network matrix *w*_*ic*_ (with dimension: number of brain regions *i* x functional networks *c)*.

Lastly, we tracked the activity of the functional networks over time (see right panel of Figure 2B). Through projection of the binarized event matrix onto the network matrix, the similarity between each functional network *c* and the whole-brain activity at each time point *b* could be assessed. This resulted in an activity matrix *A* (with the dimension: functional networks *c* x time points *b*):

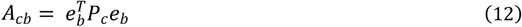

with the event matrix *e* and the projection matrix *P* The projection matrix *P* is defined as:

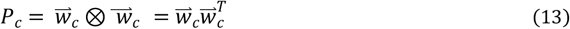

where ⊗ is the outer product operator, 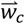 is the one of the extracted functional networks from the event matrix (the column of the matrix *w*_*c*_) and *e*_*b*_is the *b* column of the event matrix (events at time point *b*).

After retrieving the activity of each functional network over time, we calculated its probability of occurrence. We calculated the ratio of activity of each functional network in relation to overall activity (activity of all networks over time), resulting in the probability of each network *c* over time:

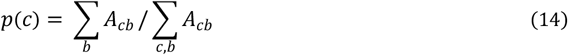

where *b* corresponds to each time point.

Using these probabilities, we computed the entropy of occurrence of each network *c*. The entropy represents the richness of switching activity between functional networks, adapted from the concept of entropy by Shannon (1948):

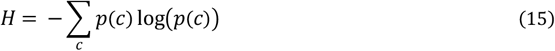

As the number of functional networks increased with higher spatial scales, we performed a normalization of the entropy. The normalization was done by dividing the entropy by the logarithm of the resulting number of networks for each spatial scale. By doing so, it was possible to compare across spatial scales. We then compared the entropy of network switching across spatial and temporal scales (see Figure 2C). We did a pairwise comparison of entropy of spatial scales using Wilcoxon tests in the empirical data and the simulated data (at the optimal temporal scale and at the temporal scale = TR).

## Results

We aimed to describe the optimal spatiotemporal scale that captured the highest information content about the temporal evolution of functional networks (as evidenced by the switching activity). We extracted time series at different parcellations at different spatial scales (from 100 to 900 regions) in the empirical data. Furthermore, we created a dynamic mean-field model to create time series at various temporal scales from milliseconds to seconds (Figure 1) and a spatial scale between 100 and 400 regions. We extracted functional networks from both simulated and empirical time series using independent component analysis. We then explored the probability of occurrence of these functional networks over time. We calculated the entropy of these probabilities’ occurrence of each network, which represents the diversity of switching activity between functional networks (Figure 2). By restricting our analysis to functional networks (as opposed to raw time series), we ensured that the information we gained on the temporal dynamics (as measured by switching activity) was relevant for whole-brain information processing.

### Agreement between empirical and simulated data

The DMF model is a neuronal model that recreates inhibitory and excitatory synaptic dynamics (including AMPA, GABA and NMDA receptors) following the structure given by the underlying anatomical connectivity. By using the steps detailed in Figure 1 and following the constraints of anatomical connectivity as provided by the structural connectome, we were able to create realistic neuronal time series at the scale of milliseconds to seconds using the DMF model. To ensure the robustness of the model, we fitted the resulting simulated BOLD time series to the empirical BOLD time series. Here, we defined a good fitting where the differences in metastability and the Kolmogorov-Smirnov statistics of the phase consistency matrices reached a minimum (see Figure S1). As can be seen in Figure S1, the fitting resulted in an optimum at a global coupling value G between 1.55 and 1.85 (depending on the spatial scale used).

In both the simulated and empirical data, some of the resulting networks resembled known classical resting state networks (see Figure 4). As our study focused on the dynamical alteration of functional networks, we aimed to ensure that the properties of the resulting functional networks from the simulation were comparable to the properties of the networks derived from the empirical time series. Therefore, we compared the number of functional networks derived from the simulated BOLD time series (see Figure S2). Here, the number of functional networks and its change across spatial scales (i.e., an increase of functional networks with increasing number of regions) were more in agreement with the empirical functional networks.

### Entropy of switching of whole-brain functional networks

The switching of whole-brain functional networks over time and their probabilities of occurrence allowed us to estimate entropy *H* as a representation of the information content of the functional network activity at various spatiotemporal scales from a probabilistic perspective. We display the entropy of spatiotemporal networks as a function of the spatial and temporal scale using empirical (Figure 3A) and simulated time series (Figure 3B). As the number of networks was contingent on the spatial scale used, we corrected the entropy for the logarithm of the number of networks to be able to compare across different spatial scales.

**Figure 3.**
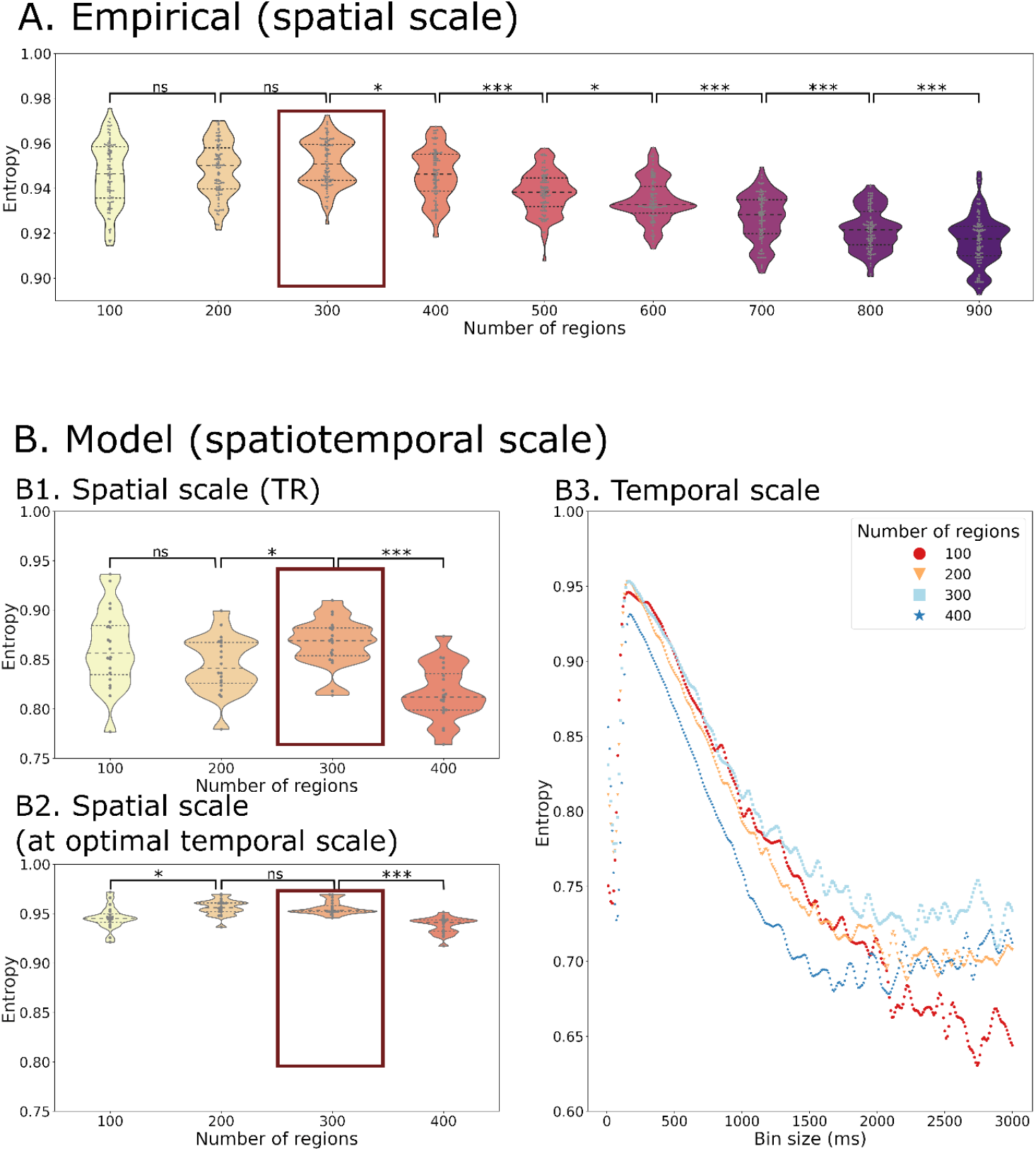
Entropy of the temporal probability of whole-brain functional networks in different spatial and temporal scales of the empirical (A) and simulated data (B). The entropy is calculated across spatial scales in the empirical data with a fixed temporal scale of 720 ms (corresponding to the TR). The simulated data gives the opportunity to explore different spatial scales at the temporal scale of the TR, 720 ms, (B1) as well as at the optimal temporal scale of 150 ms (B2). Beyond that it can be also used to explore various temporal scales and spatial scales simultaneously (B3). Both the empirical and simulated data show that the highest entropy can be found at a spatial scale of 300 regions with only a minor decrease in entropy at a spatial scale of 200 regions (marked by a red box in A and B1-B2). The highest entropy can be found at a temporal scale of 150 ms across all spatial scales (B3). Each datapoint depicts a random group of 10 subjects in the empirical data or a simulation trial simulating a group of 10 subjects. Statistical significance of comparisons between spatial scales is indicated with “ns” meaning a p-value > 0.05, * meaning < 0.05, *** meaning 0.001 (FDR-corrected).

We discovered an inverted U-shape form of the entropy *H* as a function of probability of spatiotemporal networks across time. Regarding the spatial scale, the *H* reached the highest value at a scale of 300 regions (mean simulated *H* = 0.957, mean empirical *H* = 0.951), but with only a small decrease at scales with 100 (mean simulated *H* = 0.949, mean empirical *H* = 0.946) or 400 regions (mean simulated *H* = 0.938, mean empirical *H* = 0.946). At spatial scales above 400 regions (analysis only present in empirical data, see Figure 4A), we observed a further drop in entropy (down to mean empirical *H* = 0.916 at 900 regions).

**Figure 4.**
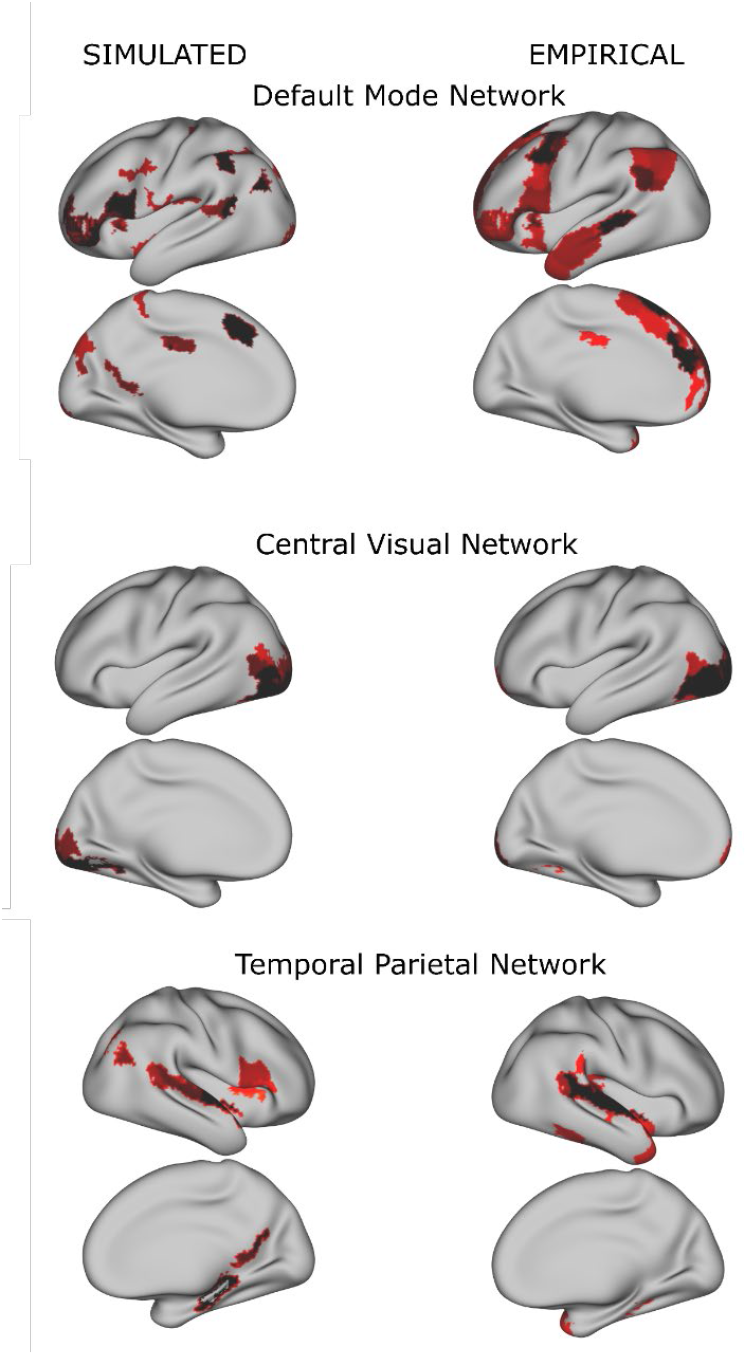
Examples of group whole-brain functional networks rendered on the standard brain. The left column has been retrieved from the simulated time series (using a TR = 720 ms), the right column from the empirical time series. Some of these networks have a high overlap with classical resting state networks (Yeo et al. 2011) such as the Default Mode Network, Central Visual Network and Temporal Parietal Network.

Beside the comparison across spatial scales, the simulated time series allowed us to compare the temporal scales (Figure 3B). Regarding the temporal scale, we found the highest entropy at an average scale of 150 ms (ranging from 140 to 160 ms, depending on the spatial scale used). Using finer or coarser temporal scales led a much greater drop in entropy (lowest value: mean simulated *H* = 0.5957) than a change of spatial scales.

Taking both spatial and temporal scales into account, the highest level of entropy could be found at a temporal scale of 150 ms and a spatial scale of 300 regions (see Figure 3 B3). The optimal temporal scale of 150 milliseconds persisted at all simulated spatial scales. Also, the effect of temporal scale on entropy was greater than the effect of spatial scale. Of note, *H* was always higher when using the empirical dataset in comparison to the simulated time series even when using the temporal scale (see Figure 3A vs. Figure 3 B1), reflecting the variability given by the empirical time series (and signals not accounted for in the dynamic mean field model).

## Discussion

In this study, we investigated the most relevant spatiotemporal scale of fundamental macroscopic dynamical processes, such as the transitions between whole-brain functional networks. We followed the temporal behaviour of functional whole-brain networks at different spatial scales and at fine-grained temporal scales from milliseconds to seconds (using a realistic whole-brain dynamic mean field model). In both empirical and simulated datasets, we generated evidence that the entropy of network switching followed an inverted U-shaped curve with a maximum at a spatial scale at about 300 regions and at a temporal scale of about 150 milliseconds. Of note, the optimal temporal scale of about 150 milliseconds persisted at all simulated spatial scales from 100 to 400 regions, indicating an absent interaction effect between spatial and temporal scales. Also, the effect of the temporal scale on entropy was much greater than the effect of spatial scale. Given the close agreement of results using simulated and empirical time series, our whole-brain network model offers an excellent opportunity to bridge analyses of brain dynamics across different neuroimaging modalities at different spatiotemporal scales, e.g. fMRI and EEG data.

Previous studies have performed comparisons between spatial scales in regard to various metrics, such as the reproducibility of resulting networks, agreement with anatomical connectivity, and prediction accuracy of neuropsychiatric conditions (Craddock et al. 2011; Arslan et al. 2018; Dadi et al. 2019; Messé 2019). However, all these studies focused on the average functional connectivity, without considering the dynamics of these networks. Only Proix et al. (2016) investigated the effect of spatial scale on the information content of brain dynamics by decomposing the time series using a principle component analysis in a whole-brain network model and found the highest eigenvalue at around 140 regions. Higher spatial scales led to an oversampling with a relative reduction of connectome density, leading to more segregated regions and an overall reduction of transmission information content across regions. Although these results are promising, they focused on separate regions rather than whole-brain networks.

Our study is the first to examine spatial and temporal scales simultaneously with a focus on brain dynamics of whole-brain networks. Given the significant evidence that maximal entropy of brain dynamics is associated with maximal transmission of information (Lungarella and Sporns 2006; Rämö et al. 2007; Shew et al. 2011; Wang et al. 2018) and is associated with cognitive performance (Niu et al. 2018; Liu et al. 2020) and consciousness (Mashour and Hudetz 2018), we chose to describe the richness of whole-brain network activity using the entropy of whole-brain network switching. Selecting the most informative spatiotemporal scale during analyses of brain dynamics can help to focus the analysis on relevant information about the dynamical behaviour of brain networks, while reducing the amount of noise (Fornito 2010), avoiding oversampling (Proix et al. 2016) and optimizing the computational cost of the analysis, i.e. removing subnetworks that are barely active and contribute little to the overall network activity.

Our findings have several implications for future research of brain dynamics. First, we were able to reproduce the finding of the optimal temporal scale of about 150 milliseconds using another dataset (Deco et al. 2019). Our findings reflect experimental results of temporal dynamics of conscious processes that operate at similar temporal scales and typically involve a rapid temporal sequence of information stabilization and transfer (Koenig et al. 2002; Van De Ville et al. 2010; Wutz et al. 2014; Salti et al. 2015; Mai et al. 2019). On top of that, our study shows that the optimal temporal scale does not depend on the spatial scale, i.e. an optimal scale of about 150 milliseconds persists across all spatial scales. For researchers aiming to extract the most relevant information content in their analyses of brain dynamics, we therefore advise to either use neuroimaging modalities operating at this optimal temporal scale (e.g. MEG or EEG) (Michel and Koenig 2018) or augment their analyses with whole-brain modeling, which allows to take other temporal scales into consideration. Second, our study provides an empirical basis for choosing the spatial scale for neuroimaging analyses with a focus on brain dynamics of whole-brain functional networks. We provided evidence that a spatial scale of about 300 regions is sufficient to capture the most relevant information on macroscopic brain dynamics. While lower scales may be associated with a loss of information, higher spatial scales introduce irrelevant and possibly more noisy functional networks. Our recommendations, based on empirical data rather than arbitrary choices, might contribute to harmonizing analyses of brain dynamics across scales.

### Limitations and outlook

There are several limitations in our methodological approach. First, we used independent component analysis to derive whole-brain functional networks at different scales. As any other higher-order statistical method, independent component analysis is not free of underlying assumptions and especially assumes maximal spatial independence of the networks (Jutten and Herault 1991). Future studies could consider additional analyses using other metrics such as network measures. However, as Arslan et al. (2018) and Hilger et al. (2020) demonstrated in their studies (Arslan et al. 2018; Hilger et al. 2020), many network measures are largely altered by the spatial scale and appropriate correction techniques should be used for such analyses across scales.

Second, our analysis was focused on the spatial scales of dynamical behaviour of whole-brain networks. Depending on the size of the networks of interest, other spatial and temporal scales might be relevant. Future studies could therefore consider exploring brain dynamics of cellular-level networks using microscale imaging tools such as optical imaging. Methods aiming at analytically bridging macro- and microscales are currently under investigation (Weiskopf et al. 2015; Larivière et al. 2019; Gao et al. 2020).

Third, both the estimation of the whole-brain functional networks as well as the calculation of the entropy of the network switching activity was done on a group level. Comparing the entropy of network switching on an individual level would allow to relate individual cognition to dynamical behaviour of brain networks.

Overall, our results suggest that whole-brain functional brain networks operate at an optimum of about 300 regions and a timescale of about 150 milliseconds. We contribute to the understanding of the dynamical behaviour of whole-brain networks, which could inspire future human neuroimaging studies to harmonize spatiotemporal scales and use dynamical models to create connections between micro- and macroscopic scales.

## Funding

This work was supported by a short-term fellowship of the European Molecular Biology Organization (n. 7366) and by an InEurope fellowship of the International Brain Research Organization of XK. AL is supported by Swiss National Science Foundation Sinergia grant no. 170873. GD is supported by AWAKENING Using whole-brain models perturbational approaches for predicting external stimulation to force transitions between different brain states (ref. PID2019-105772GB-I00, AEI FEDER EU), funded by the Spanish Ministry of Science, Innovation and Universities (MCIU), State Research Agency (AEI) and European Regional Development Funds (FEDER), by HBP SGA3 Human Brain Project Specific Grant Agreement 3 (Grant Agreement No. 945539), funded by the EU H2020 FET Flagship program and by SGR Research Support Group support (ref. 2017 SGR 1545), funded by the Catalan Agency for Management of University and Research Grants (AGAUR). MLK is supported by the ERC Consolidator Grant: CAREGIVING (n. 615539), Center for Music in the Brain, funded by the Danish National Research Foundation (DNRF117), and Centre for Eudaimonia and Human Flourishing funded by the Pettit and Carlsberg Foundations.

## Notes

### Competing Interest Statement

The authors have declared no competing interest.

## References

Achard S, Salvador R, Whitcher B, Suckling J, Bullmore E. 2006. A resilient, low-frequency, small-world human brain functional network with highly connected association cortical hubs. J Neurosci Off J Soc Neurosci. 26:63–72.

Alexandrov YI. 1999. Physiological Regularities of the Dynamics of Individual Experience and the “Stream of Consciousness”, in Neural Bases and Psychological Aspects of Consciousness, Teddei-Ferretti, C., and Musio, C., Eds., Singapore: World Scientific, 1999, p. 201. In: Neural Bases and Psychological Aspects of Consciousness. Singapore: World Scientific. p. 201.

Arslan S, Ktena SI, Makropoulos A, Robinson EC, Rueckert D, Parisot S. 2018. Human brain mapping: A systematic comparison of parcellation methods for the human cerebral cortex. NeuroImage. 170:5–30.

Ashburner J. 2007. A fast diffeomorphic image registration algorithm. NeuroImage. 38:95–113.

Cornblath EJ, Ashourvan A, Kim JZ, Betzel RF, Ciric R, Adebimpe A, Baum GL, He X, Ruparel K, Moore TM, Gur RC, Gur RE, Shinohara RT, Roalf DR, Satterthwaite TD, Bassett DS. 2020. Temporal sequences of brain activity at rest are constrained by white matter structure and modulated by cognitive demands. Commun Biol. 3:1–12.

Craddock RC, James GA, Holtzheimer PE, Hu XP, Mayberg HS. 2011. A whole brain fMRI atlas generated via spatially constrained spectral clustering. Hum Brain Mapp. 33:1914–1928.

Dadi K, Rahim M, Abraham A, Chyzhyk D, Milham M, Thirion B, Varoquaux G, Alzheimer’s Disease Neuroimaging Initiative. 2019. Benchmarking functional connectome-based predictive models for resting-state fMRI. NeuroImage. 192:115–134.

Deco G, Cruzat J, Kringelbach ML. 2019. Brain songs framework used for discovering the relevant timescale of the human brain. Nat Commun. 10:583.

Deco G, Kringelbach ML, Jirsa VK, Ritter P. 2017. The dynamics of resting fluctuations in the brain: metastability and its dynamical cortical core. Sci Rep. 7:3095.

Deco G, Ponce-Alvarez A, Hagmann P, Romani GL, Mantini D, Corbetta M. 2014. How Local Excitation–Inhibition Ratio Impacts the Whole Brain Dynamics. J Neurosci. 34:7886– 7898.

Deco G, Ponce-Alvarez A, Mantini D, Romani GL, Hagmann P, Corbetta M. 2013. Resting-state functional connectivity emerges from structurally and dynamically shaped slow linear fluctuations. J Neurosci Off J Soc Neurosci. 33:11239–11252.

Engel AK, Fries P, Singer W. 2001. Dynamic predictions: oscillations and synchrony in top-down processing. Nat Rev Neurosci. 2:704–716.

Fornito. 2010. Network scaling effects in graph analytic studies of human resting-state fMRI data. Front Syst Neurosci.

Gao R, van den Brink RL, Pfeffer T, Voytek B. 2020. Neuronal timescales are functionally dynamic and shaped by cortical microarchitecture. eLife. 9:e61277.

Glasser MF, Sotiropoulos SN, Wilson JA, Coalson TS, Fischl B, Andersson JL, Xu J, Jbabdi S, Webster M, Polimeni JR, Van Essen DC, Jenkinson M, WU-Minn HCP Consortium. 2013. The minimal preprocessing pipelines for the Human Connectome Project. NeuroImage. 80:105–124.

Glerean E, Salmi J, Lahnakoski JM, Jääskeläinen IP, Sams M. 2012. Functional magnetic resonance imaging phase synchronization as a measure of dynamic functional connectivity. Brain Connect. 2:91–101.

Hilger K, Fukushima M, Sporns O, Fiebach CJ. 2020. Temporal stability of functional brain modules associated with human intelligence. Hum Brain Mapp. 41:362–372.

Horn A, Blankenburg F. 2016. Toward a standardized structural–functional group connectome in MNI space. NeuroImage. 124:310–322.

Horn A, Li N, Dembek TA, Kappel A, Boulay C, Ewert S, Tietze A, Husch A, Perera T, Neumann W-J, Reisert M, Si H, Oostenveld R, Rorden C, Yeh F-C, Fang Q, Herrington TM, Vorwerk J, Kühn AA. 2018. Lead-DBS v2: Towards a comprehensive pipeline for deep brain stimulation imaging. NeuroImage. 184:293–316.

Horn A, Reich M, Vorwerk J, Li N, Wenzel G, Fang Q, Schmitz-Hübsch T, Nickl R, Kupsch A, Volkmann J, Kühn AA, Fox MD. 2017. Connectivity Predicts Deep Brain Stimulation Outcome in Parkinson Disease. Ann Neurol. 82:67–78.

Jutten C, Herault J. 1991. Blind separation of sources, part I: An adaptive algorithm based on neuromimetic architecture. Signal Process. 24:1–10.

Koenig T, Prichep L, Lehmann D, Sosa PV, Braeker E, Kleinlogel H, Isenhart R, John ER. 2002. Millisecond by Millisecond, Year by Year: Normative EEG Microstates and Developmental Stages. NeuroImage. 16:41–48.

Kreher BW, Mader I, Kiselev VG. 2008. Gibbs tracking: a novel approach for the reconstruction of neuronal pathways. Magn Reson Med. 60:953–963.

Larivière S, Vos de Wael R, Paquola C, Hong S-J, Mišić B, Bernasconi N, Bernasconi A, Bonilha L, Bernhardt BC. 2019. Microstructure-Informed Connectomics: Enriching Large-Scale Descriptions of Healthy and Diseased Brains. Brain Connect. 9:113–127.

Liégeois R, Li J, Kong R, Orban C, Van De Ville D, Ge T, Sabuncu MR, Yeo BTT. 2019. Resting brain dynamics at different timescales capture distinct aspects of human behavior. Nat Commun. 10:2317.

Liu M, Liu X, Hildebrandt A, Zhou C. 2020. Individual Cortical Entropy Profile: Test–Retest Reliability, Predictive Power for Cognitive Ability, and Neuroanatomical Foundation. Cereb Cortex Commun. 1.

Lopes-dos-Santos V, Ribeiro S, Tort ABL. 2013. Detecting cell assemblies in large neuronal populations. J Neurosci Methods. 220:149–166.

Lungarella M, Sporns O. 2006. Mapping Information Flow in Sensorimotor Networks. PLOS Comput Biol. 2:e144.

Lurie DJ, Kessler D, Bassett DS, Betzel RF, Breakspear M, Kheilholz S, Kucyi A, Liégeois R, Lindquist MA, McIntosh AR, Poldrack RA, Shine JM, Thompson WH, Bielczyk NZ, Douw L, Kraft D, Miller RL, Muthuraman M, Pasquini L, Razi A, Vidaurre D, Xie H, Calhoun VD. 2020. Questions and controversies in the study of time-varying functional connectivity in resting fMRI. Netw Neurosci. 4:30–69.

Mai A-T, Grootswagers T, Carlson TA. 2019. In search of consciousness: Examining the temporal dynamics of conscious visual perception using MEG time-series data. Neuropsychologia. 129:310–317.

Marčenko VA, Pastur L. 1967. Distribution of eigenvalues for some sets of random matrices. Math USSR Sb. 1:457–483.

Mashour GA, Hudetz AG. 2018. Neural Correlates of Unconsciousness in Large-Scale Brain Networks. Trends Neurosci. 41:150–160.

Meer JN van der, Breakspear M, Chang LJ, Sonkusare S, Cocchi L. 2020. Movie viewing elicits rich and reliable brain state dynamics. Nat Commun. 11:5004.

Messé A. 2019. Parcellation influence on the connectivity-based structure–function relationship in the human brain. Hum Brain Mapp. 41:1167–1180.

Michel CM, Koenig T. 2018. EEG microstates as a tool for studying the temporal dynamics of whole-brain neuronal networks: A review. NeuroImage, Brain Connectivity Dynamics. 180:577–593.

Niu Y, Wang B, Zhou M, Xue J, Shapour H, Cao R, Cui X, Wu J, Xiang J. 2018. Dynamic Complexity of Spontaneous BOLD Activity in Alzheimer’s Disease and Mild Cognitive Impairment Using Multiscale Entropy Analysis. Front Neurosci. 12.

Peyrache A, Benchenane K, Khamassi M, Wiener SI, Battaglia FP. 2010. Principal component analysis of ensemble recordings reveals cell assemblies at high temporal resolution. J Comput Neurosci. 29:309–325.

Proix T, Spiegler A, Schirner M, Rothmeier S, Ritter P, Jirsa VK. 2016. How do parcellation size and short-range connectivity affect dynamics in large-scale brain network models? NeuroImage. 142:135–149.

Rämö P, Kauffman S, Kesseli J, Yli-Harja O. 2007. Measures for information propagation in Boolean networks. Phys Nonlinear Phenom. 227:100–104.

Saenger VM, Kahan J, Foltynie T, Friston K, Aziz TZ, Green AL, van Hartevelt TJ, Cabral J, Stevner ABA, Fernandes HM, Mancini L, Thornton J, Yousry T, Limousin P, Zrinzo L, Hariz M, Marques P, Sousa N, Kringelbach ML, Deco G. 2017. Uncovering the underlying mechanisms and whole-brain dynamics of deep brain stimulation for Parkinson’s disease. Sci Rep. 7:9882.

Salti M, Monto S, Charles L, King J-R, Parkkonen L, Dehaene S. 2015. Distinct cortical codes and temporal dynamics for conscious and unconscious percepts. eLife. 4.

Schaefer A, Kong R, Gordon EM, Laumann TO, Zuo X-N, Holmes AJ, Eickhoff SB, Yeo BTT. 2018. Local-Global Parcellation of the Human Cerebral Cortex from Intrinsic Functional Connectivity MRI. Cereb Cortex. 28:3095–3114.

Setsompop K, Kimmlingen R, Eberlein E, Witzel T, Cohen-Adad J, McNab JA, Keil B, Tisdall MD, Hoecht P, Dietz P, Cauley SF, Tountcheva V, Matschl V, Lenz VH, Heberlein K, Potthast A, Thein H, Van Horn J, Toga A, Schmitt F, Lehne D, Rosen BR, Wedeen V, Wald LL. 2013. Pushing the limits of in vivo diffusion MRI for the Human Connectome Project. NeuroImage. 80:220–233.

Shannon CE. 1948. A Mathematical Theory of Communication. Bell Syst Tech J. 27:379–423.

Shew WL, Yang H, Yu S, Roy R, Plenz D. 2011. Information capacity and transmission are maximized in balanced cortical networks with neuronal avalanches. J Neurosci Off J Soc Neurosci. 31:55–63.

Stephan KE, Weiskopf N, Drysdale PM, Robinson PA, Friston KJ. 2007. Comparing hemodynamic models with DCM. NeuroImage. 38:387–401.

Stitt I, Hollensteiner KJ, Galindo-Leon E, Pieper F, Fiedler E, Stieglitz T, Engler G, Nolte G, Engel AK. 2017. Dynamic reconfiguration of cortical functional connectivity across brain states. Sci Rep. 7:1–14.

Tagliazucchi E, Balenzuela P, Fraiman D, Chialvo DR. 2012. Criticality in large-scale brain FMRI dynamics unveiled by a novel point process analysis. Front Physiol. 3:15.

Tang Y-Y, Rothbart MK, Posner MI. 2012. Neural correlates of establishing, maintaining, and switching brain states. Trends Cogn Sci. 16:330–337.

Thompson GJ, Magnuson ME, Merritt MD, Schwarb H, Pan W-J, McKinley A, Tripp LD, Schumacher EH, Keilholz SD. 2013. Short-time windows of correlation between large-scale functional brain networks predict vigilance intraindividually and interindividually. Hum Brain Mapp. 34:3280–3298.

Van De Ville D, Britz J, Michel CM. 2010. EEG microstate sequences in healthy humans at rest reveal scale-free dynamics. Proc Natl Acad Sci. 107:18179–18184.

Van Essen DC, Smith SM, Barch DM, Behrens TEJ, Yacoub E, Ugurbil K. 2013. The WU-Minn Human Connectome Project: An overview. NeuroImage, Mapping the Connectome. 80:62–79.

Vidaurre D, Smith SM, Woolrich MW. 2017. Brain network dynamics are hierarchically organized in time. Proc Natl Acad Sci. 114:12827–12832.

Wang DJJ, Jann K, Fan C, Qiao Y, Zang Y-F, Lu H, Yang Y. 2018. Neurophysiological Basis of Multi-Scale Entropy of Brain Complexity and Its Relationship With Functional Connectivity. Front Neurosci. 12.

Weiskopf N, Mohammadi S, Lutti A, Callaghan MF. 2015. Advances in MRI-based computational neuroanatomy: from morphometry to in-vivo histology. Curr Opin Neurol. 28:313–322.

Wildie M, Shanahan M. 2012. Metastability and chimera states in modular delay and pulse-coupled oscillator networks. Chaos Interdiscip J Nonlinear Sci. 22:043131.

Wong K-F. 2006. A Recurrent Network Mechanism of Time Integration in Perceptual Decisions. J Neurosci. 26:1314–1328.

Wutz A, Weisz N, Braun C, Melcher D. 2014. Temporal Windows in Visual Processing: “Prestimulus Brain State” and “Poststimulus Phase Reset” Segregate Visual Transients on Different Temporal Scales. J Neurosci. 34:1554–1565.

Yeo BT, Krienen FM, Sepulcre J, Sabuncu MR, Lashkari D, Hollinshead M, Roffman JL, Smoller JW, Zöllei L, Polimeni JR, Fischl B, Liu H, Buckner RL. 2011. The organization of the human cerebral cortex estimated by intrinsic functional connectivity. J Neurophysiol. 106:1125–1165.

Yoo HB, Moya BE, Filbey FM. 2020. Dynamic functional connectivity between nucleus accumbens and the central executive network relates to chronic cannabis use. Hum Brain Mapp. 41:3637–3654.

Yuan J, Li X, Zhang J, Luo L, Dong Q, Lv J, Zhao Y, Jiang X, Zhang S, Zhang W, Liu T. 2018. Spatio-temporal modeling of connectome-scale brain network interactions via time-evolving graphs. NeuroImage. 180:350–369.

